# Enhancing Predictive Accuracy in Immunotherapy Models through Data Integration and Parameter Identifiability

**DOI:** 10.64898/2026.01.12.699041

**Authors:** Yixuan Wang, Daniel R. Bergman, Erica Trujillo, Andrea Ziblat, Anthony A. Fernald, Lie Li, Alexander T. Pearson, Randy F. Sweis, Trachette L. Jackson

**Affiliations:** Department of Mathematics, University of Michigan, Ann Arbor, MI 48109, USA; Department of Medicine, Section of Hematology/Oncology, The University of Chicago, Chicago, IL 60637, USA

## Abstract

Immune checkpoint inhibitors (ICIs), a class of immunotherapy, offer promising benefits but face challenges such as low response rates to be used as a broadly effective treatment for all patients. In this study, we use a set of ordinary different equation (ODE) models and bladder cancer *in vivo* data as a case study to outline a biologically informed, data-driven framework for formulating, calibrating and validating immunotherapy models, and thus ensuring their predictive reliability. We consider multiple treatment scenarios and distinct immune cell-mediated killing mechanisms for tumor cells of different antigenicity. By integrating sensitivity analysis and identifiability analysis with targeted experimental design, we demonstrate how mathematical models can move beyond qualitative insight to quantitative prediction. We generate virtual cohorts to show that insufficient data integration leads to systematically overestimated therapeutic benefits of ICIs. We also explore dosing schedules that enhance survival or reduce dosage without compromising survival.

## 1 Introduction

Cancer remains a leading cause of morbidity and mortality worldwide, posing a persistent challenge despite significant advances in diagnosis and treatment. Among the many forms of cancer, bladder cancer is one of the most prevalent malignancies globally, with approximately 550,000 new cases and 200,000 deaths annually [31, 44]. In the United States alone, over 80,000 new cases and 17,000 deaths occur each year [48]. While the five-year survival rate for bladder cancer in general is around 80%, metastatic bladder cancer has a dismal median survival time of shorter than 32 months even with modern combination immunotherapies [41]. Therefore, the unmet need for effective advanced and metastatic bladder cancer treatments has sparked significant research interest and clinical initiatives.

Historically, chemotherapy has been the cornerstone of bladder cancer treatment. Nonetheless, chemotherapy as a first-line management strategy has been limited in improving survival for advanced cases of bladder cancer [33]. Recent advances in immunotherapy, targeted therapy, and antibody-drug conjugates have provided additional lines of treatment for patients who respond poorly to chemotherapy or are ineligible for it [31]. Combination immunotherapy with pembrolizumab plus antibody drug conjugate therapy with enfortumab vedotin has now emerged as the standard of care in the front line. Among the various immunotherapeutic approaches — such as monoclonal antibodies, oncolytic viruses, adoptive cell therapy, and cancer vaccines — immune checkpoint inhibitors (ICIs) have emerged as a particularly promising class of treatments.

Unlike traditional chemotherapy, which indiscriminately kills or slows the growth of rapidly dividing cells [13, 38], immunotherapy stimulates or enhances the immune system’s ability to recognize and destroy cancer cells [50, 46]. ICIs block inhibitory immunoreceptors, like those associated with the programmed death-1/programmed death-ligand 1 (PD-1/PD-L1) checkpoint, which plays a key role in tumor immune evasion [23, 18]. The FDA has approved multiple anti-PD-1 or anti-PD-L1 [43] ICIs as second-line treatments for bladder cancer [43]. ICIs as first-line therapy have been granted conditional approval or are in clinical trials [43]. Research showed ICIs to have advantages in durability and improvement of survival over chemotherapy or targeted therapies [21], but the benefits of ICIs also proved to be more complex and uncertain than traditional anti-tumor therapies. Despite their success, ICIs face challenges such as low response rates in many patients [34, 51, 17]. Thus, there has been a great focus in preclinical and clinical research on improving ICI efficacy. Given the urgent need for better therapeutic strategies, bladder cancer serves as an ideal case study for developing a systematic way to predict ICI treatment outcomes and optimize therapeutic strategies for individual patients.

Mathematical and computational modeling has become essential for understanding and optimizing immunotherapy. Computational models can provide valuable insights into tumor-immune dynamics, enabling predictions of therapeutic responses and informing treatment strategies. Common model types include differential ordinary or partial differential equation-based models (ODE or PDE), evolutionary game theory models, discrete-time models, Markov chain, agent-based models (ABM) [53], and machine learning models. In bladder cancer, computational models have been used to predict responses to BCG immunotherapy [5, 42], synergize immunotherapy with radiotherapy [47], develop new biomarkers to predict ICI efficacy [58], and describe tumor evolution [6]. In particular, ODE and PDE-based models have been extensively used to model tumor-immune interactions with ICI monotherapy or combination therapy [37, 26, 27, 30, 29], understand pharmacokinetic or pharmacodynamic (PK/PD) properties of anti-PD1 to help with dose selection [12] and predict responses to chemotherapy or anti-PD-L1 ICI [14]. We previously developed an ODE model and an ABM to predict tumor responses to ICI monotherapy [56, 55] and ICI in combination with FGFR3 targeted therapy [2].

Nevertheless, modeling biological systems presents significant challenges, particularly in parameter estimation and model validation. For example, Brady and Enderling highlighted the importance of calibration and validation in producing models that give reliable and clinically relevant predictions of novel cancer treatments [3]. Model calibration is more than finding parameters that result in simulations that fit experimental data. Two key concepts in this context are structural identifiability and practical identifiability. Structural identifiability determines whether a model’s parameters can be uniquely estimated from idealized, noise-free data [9, 45, 54]. In contrast, practical identifiability considers whether parameters can be reliably inferred from real-world, noisy experimental data [40, 11]. Ensuring parameter identifiability is critical for producing trustworthy predictions and guiding treatment optimizations.

To address the these challenges, we present a data-driven framework for evaluating and enhancing parameter identifiability in immunotherapy models. By developing biologically informed mathematical models that simulate the effects of ICI therapy in bladder cancer, we demonstrate that integrative analyses across multiple *in vivo* datasets improve the reliability and clinical relevance of predictions for ICI treatment outcomes. Using three *in vivo* bladder cancer datasets, we delineate how to assess and enhance the identifiability of crucial immune-related parameters given different types and amount of data. We show that insufficient data integration can lead to overestimated tumor reduction and inaccurate survival predictions. Our study also compares therapeutic scenarios, including complete immune checkpoint blockade, clinically relevant anti-PD-L1 dosing regimens, and alternative dosing schedules. We examine the optimal treatment strategies to enhance survival of the virtual cohort or to reduce dosage without compromising survival. Additionally, we investigate how tumor antigenicity influences the efficacy of ICIs. Collectively, these findings underscore the critical role of data integration and parameter identifiability in enhancing the predictive accuracy of cancer immunotherapy models, thereby strengthening their utility for clinical decision-making.

## 2 Methods

### 2.1 Description of experiments

We use three sets of *in vivo* data from mouse bladder cancer studies. Table 1 compares the immunocompetency of mice, antigenicity of tumor cells, number of mice, treatment conditions and type of measurements in each experiment. Six to eight-week-old female RAG1 KO or C57BL/6J mice were obtained from The Jackson Laboratory. Mice were housed in a specific pathogen-free animal facility at the University of Chicago and used in accordance with the animal experimental guidelines set by the University of Chicago Animal Care and Use Committee (IACUC). All experimental animal procedures were approved by the IACUC. All mice were sacrificed in accordance with IACUC guidelines for humane endpoints. In all three data sets, 1 × 10^6^ MB49 cells, a model derived from a chemical carcinogen-induced urothelial carcinoma, were subcutaneously injected in mice. Each data set has different experimental conditions, types, and measurements that are timed differently.

**Table 1:** Comparison of all *in vivo* data sets. C57BL/6J mice are immunocompetent. RAG1 KO mice are immunocompromised. Treatment in Data Set 1 is with an anti-PD-L1 antibody.

|  | Data Set 1 [39] | Data Set 2 [55] | Data Set 3 |
| --- | --- | --- | --- |
| Type of mice used | C57BL/6J | C57BL/6J,<br>RAG1 | C57BL/6J |
| Antigen effect shown | No | No | Yes |
| No-treatment group | Yes | Yes | Yes |
| Treatment group | Yes | No | No |
| Types of measurement | Tumor volume | Tumor volume,<br>CTL density | Tumor volume |

The first data set contains tumor volume measurements in immunocompetent (C57BL/6J) mice with (n = 5) or without (n = 5) anti-PD-L1 treatments following a realistic dosing regimen [39]. We denote it “Data Set 1”. Tumor volumes were measured on days 10, 14, 19, 21, and 25, with day 25 being the end point. Anti-PD-L1 antibody therapy started on day 7 when the tumor was first palpable. Mice were randomly assigned to intraperitoneal injection of phosphate buffered saline (PBS control) or anti-PD-L1 therapy (clone 10F.9G2; BioXcell). From day 7, 100 µg of anti-PD-L1 antibody is administered every three days except on the weekends, with the last dose on day 24. The anti-PDL1 dosing scheme is chosen based on preclinical dosing of antibodies targeting the PD-L1 immune checkpoint where patients are treated with anti-PD-L1 antibodies in an intermittent schedule given the prolonged half-life of immune checkpoint antibodies [15, 52].

The second data set contains tumor volume measurements in immunocompromised (RAG1 KO, n = 27) and immunocompetent (C57BL/6J, n = 24) mice, as well as endpoint intra-tumoral CTL density measurements in C57BL/6J mice [55]. Four types of MB49 cells with different expression levels of the model antigen SIY (SIYRYYGL) were used: Zs green (no SIY), L14 (low SIY), H1 (high SIY), and a mix of L14 and H1 cells with 1:1 ratio. Each type of MB49 cell was injected into five to seven mice of each strain. Tumor volumes were measured on days 7, 10, 12, 14, 17, and 19. Intratumoral CTL density was measured at the endpoint (Day 20). We denote this “Data Set 2”. In the third data set, control MB49 cells and MB49 cells with different SIY antigen expression levels were subcutaneously injected in five C57BL/6J mice, respectively. The level of SIY expression was determined by Fluorescence-Activated Cell Sorting (FACS), with SIY 1x denoting intermediate expression and SIY 2x denoting the highest expression. We denote this as “Data Set 3”. Tumor volumes are measured on days 7, 10, 13, 16, 19, 22, 25, 27, and 29, with day 29 being the endpoint. Since tumor volumes of the control and SIY 1x cell lines do not show significant separation and it is easier to directly compare the antigenicity of the cell lines with SIY antigens, we only use data from the SIY 1x and 2x experiments. We refer to SIY 1x tumor cells as low antigen (LA) tumor cells and SIY 2x tumor cells as high antigen (HA) tumor cells.

### 2.2 Mathematical models

We use ordinary differential equations (ODEs) to model the tumor-immune dynamics with different levels of immune checkpoint activity: active, blocked or partially blocked. In most experiments analyzed in this study, mice were subcutaneously injected with a single tumor cell type. The only exception is the mix of L14 and H1 cells in Data Set 2. However, tumor growth in Data Set 2 does not show meaningful separation across cell lines in both RAG1 KO and C57BL/6J mice. Therefore, to use the *in vivo* data to calibrate and validate our models, we only consider one type of tumor cell in our ODE model. Studies showed that CTLs execute their cell-killing function through at least two different mechanisms [7, 32]. The first process is perforin/granzyme-mediated, and happens faster than the second process, which is through the Fas/FasL pathway [1, 7, 57, 19]. Evidence also indicated that the switch from fast to slow killing is associated with the a reduction in antigen presence [35]. We were the first to mathematically model two types of immune-mediated tumor cell killing [56, 55], and we extend and apply this modeling approach in the current study.

Equation 1 shows the one-phenotype model with an active immune checkpoint. This is also referred to as the no-treatment or control scenario.

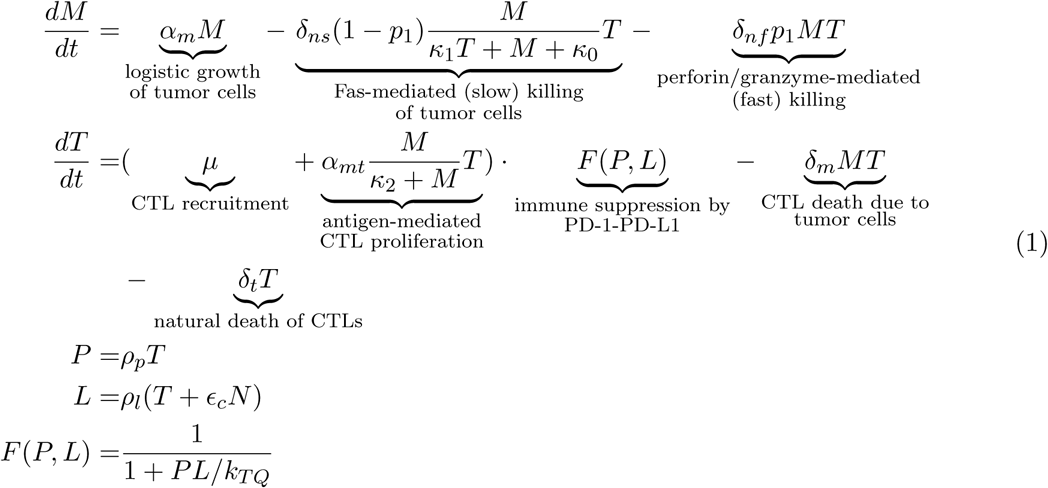

The model variables and parameters are described in Tables 2 and 1.

**Table 2:**
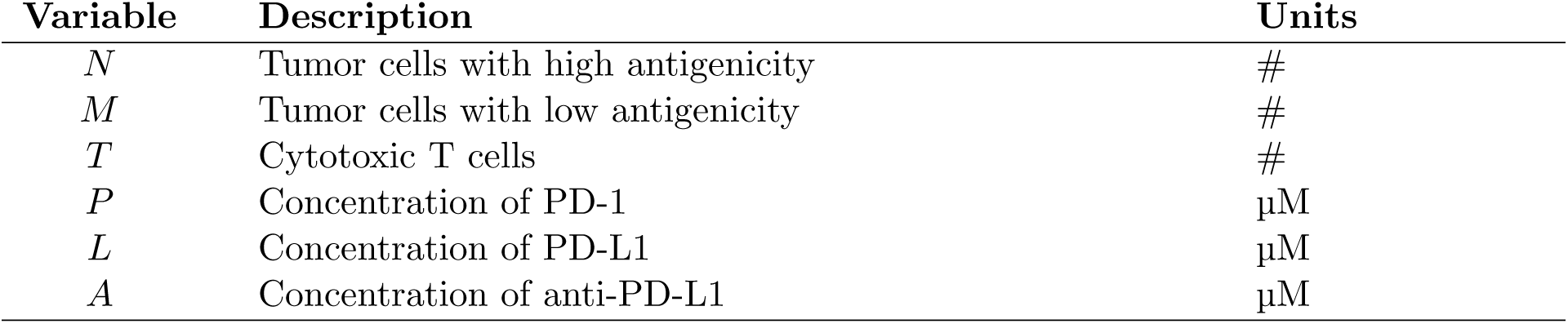
Variables for the ODE models. We use the conversion factor 10^6^ tumor cells equal 1mm^3^[**10]**.

| Variable | Description | Units |
| --- | --- | --- |
| $N$ | Tumor cells with high antigenicity | # |
| $M$ | Tumor cells with low antigenicity | # |
| $T$ | Cytotoxic T cells | # |
| $P$ | Concentration of PD-1 | $\mu\text{M}$ |
| $L$ | Concentration of PD-L1 | $\mu\text{M}$ |
| $A$ | Concentration of anti-PD-L1 | $\mu\text{M}$ |

We consider two treatment scenarios of immune checkpoint therapy: complete checkpoint blockade and anti-PD-L1 treatment following the dosing schedule used experimentally. The first scenario assumes that the ICIs are 100% effective and models the best possible treatment results. In this case, the immunosuppressive effect of PD-1/PD-L1 complexes in Equation 1 is eliminated, and *F* (*P, L*) = 1. The second scenario aligns with the treatment administered to mice in Data Set 1 and is thus more realistic. Starting from day 7, 100 µg of anti-PD-L1 antibody is administered every three days except on the weekends. In this case, the immunosuppressive effects of the PD-1/PD-L1 complexes are modulated but not completely avoided. Following the derivation in [39], we model the post-treatment immunosuppressive effect as

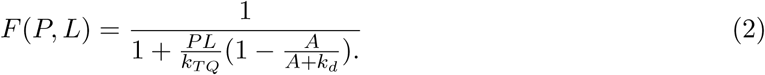

The depletion of the treatment drug anti-PD-L1 follows the following equation:

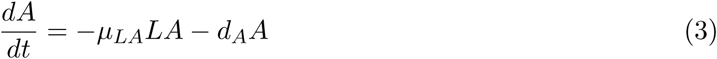

where *µ_LA_* is the depletion rate of aPD-L1 and *δ_A_*is the natural degradation rate of aPD-L1. This model contains more parameters and thus poses more challenges to parameter estimation than the treatment scenario assuming complete checkpoint blockade.

### 2.3 Structural identifiability

Structural identifiability assesses whether it is theoretically possible to uniquely determine model parameters under the assumption of perfect data—that is, data that are noise-free and continuous in both time and space. We use the “genSSI (Generating Series for testing Structural Identifiability) 2.0” software toolbox implemented in MATLAB to analyze the structural identifiability of the ODE models. In general, GenSSI can handle systems represented by a set of linear/non-linear differential equations of the form given below:

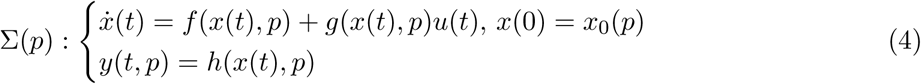

where *x* ∈ R*^n^* is the *n*-dimensional state variable, *u* ∈ R*^m^* a *m*-dimensional control, *y* ∈ R*^r^* is the *r*-dimensional observation of the state varia bles, and *x*_0_(*p*) are the initial conditions that may depend on the parameters. The unknown model parameters are *p* ∈ **P** (**P** ⊆ R*^q^*). Functions *f, g, h* are assumed to be analytic. [8] In our case, *u* = 0, and *g* = 0.

A parameter *p* = (*p*_1_*, p*_2_*, …, p_n_* ) is structurally globally identifiable if for almost any **p**^∗^ ∈ **P**,

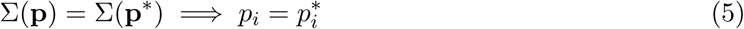

A parameter *p* is structurally locally identifiable if for almost any **p**^∗^ ∈ **P**, there exists a neighborhood **V**(**p**^∗^) such that

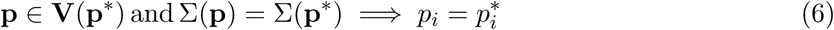

Otherwise, the parameter is structurally non-identifiable. [9] See Section S6.2.1 in the supplemental material for additional details.

We start by analyzing the ODE model with a completely blocked checkpoint (Equation 1 with *F* (*P, L*) = 1) due to its simpler formulation with fewer parameters compared to the model with an active immune checkpoint. This model is not structurally globally identifiable because the Jacobian of the exhaustive summary is rank deficient, which means that we cannot uniquely determine the model parameters even in the most ideal scenario. Without performing new experiments, we have to reduce the number of parameters to make this model globally structurally identifiable. We first combine the probability and rate of the two immune cell kill mechanisms into one parameter for each.

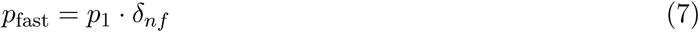

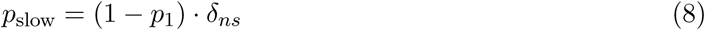

For notation purposes, we refer to *p*_fast_ as the “rate of fast killing” and *p*_slow_ as the “rate of slow killing”. Then we fix *κ*_0_*, κ*_1_ and *κ*_2_ at the values shown in Table 1. The model becomes globally identifiable as the Jacobian of the exhaustive summary is rank complete. The identifiability tableau is shown in Figure S1. Specifically, these results imply that now we can potentially uniquely estimate the CTL recruitment rate (*µ*), the rate of rapid killing (*p*_fast_), and the tumor proliferation rate (*α_m_*) from experimental data. These parameters are the top three most sensitive parameters for the model with complete checkpoint blockade in terms of their partial rank correlation coefficient (PRCC) with respect to endpoint tumor volume according to our global sensitivity analysis in [56]. They are also essential to our study of the differentiated immune activity toward tumor cells with different antigen loads.

The ODE model for tumor-immune dynamics with an active immune checkpoint in Equation 1 is also not structurally identifiable because the Jacobian of the exhaustive summary is rank deficient. In addition to fixing *κ*_0_*, κ*_1_ and *κ*_2_, we also have to fix parameters related to the PD-1/PD-L1 checkpoint, namely *ρ_p_, ρ_l_* and *ɛ_c_* to make the model structurally globally identifiable. The identifiability tableau is shown in Figure S1, the same as the one corresponding to the checkpoint blocked model. Note that this does not mean the Jacobian matrices of the checkpoint blocked and checkpoint active models are the same as the identifiability tableau only shows whether an entry of the Jacobian matrix is zero or not. The choice of fixing these parameters is partly due to global sensitivity analysis of model parameters following the method outlined in [56], and partly due to the research objective of out models. The most sensitive parameters for the ODE model with an active immune checkpoint are by far *α_m_*, followed by *p*_fast_ and *µ*. The PRCCs of other parameters are much smaller in magnitude, meaning that fixing them has a relatively small impact on the model output. Since *µ* and *p*_fast_ are the most sensitive parameters in terms of end point tumor volume after checkpoint blockade therapy, it is crucial to estimate *µ* and *p*_fast_ with confidence to accurately predict responses to checkpoint blockade therapy. Moreover, the focus of our study is to model distinct mechanisms of immune responses to tumors cells, not the expression of PD-1 or PD-L1 on tumor and immune cells.

### 2.4 Practical identifiability

Complete and noise-free time series data required for structural identifiability are rarely available in practice. Therefore, we perform a practical identifiability analysis to assess whether parameters can be reliably estimated from our experimental data sets. Since structural identifiability is a necessary condition for practical identifiability, we use equations 1, 2 and 3 with the modifications mentioned in Section 2.3 from this point onwards. Suppose *p* represents model parameters and *z* represents observed data. The goal is to estimate the posterior distribution *P* (*p*|*z*). For the no-treatment model with an active immune checkpoint, we focus on the practical identifiability of *α_m_, µ* and *p*_fast_ for reasons mentioned in Section 2.3. The treatment model introduces three more parameters *d_a_, k_d_*and *µ_la_*. Based on our sensitivity analysis of the model, only *d_a_* and *k_d_* have significant, positive PRCC values. Therefore, for the treatment model, in addition to *α_m_, µ* and *p*_fast_, we also evaluate the practical identifiability of *d_a_* and *k_d_*. Specifically, we evaluate whether each parameter yields a posterior distribution with a well-defined mode. A unimodal posterior distribution indicates practical identifiability. Otherwise, the parameter is practically unidentifiable. We use the Markov chain Monte Carlo (MCMC) method using Metropolis–Hastings sampling [20] to conduct this analysis. See Section S6.2.2 of the supplemental material for more details.

### 2.5 Virtual cohort generation

Virtual cohorts of tumors with diverse tumor-immune characteristics are often used to simulate population’s varied responses to immunotherapy treatments. There are various ways to create such virtual cohorts depending on the availability of data. Without *in vivo* data, modelers can create a virtual cohort by randomly sampling the prior distributions of parameters, which are based on their knowledge and assumptions of the parameters before observing any experimental data. The prior distributions are commonly assumed to be uniform. With *in vivo* data, one can create a virtual cohort from data-driven posterior distributions obtained from practical identifiability analysis. We use both approaches to generate virtual cohorts using Data set 1 and use the second approach for Data sets 2 and 3.

We first use Dataset 1 to examine how varying degrees of data integration influence practical identifiability and model predictions. We construct three virtual cohorts using i) no data, ii) no-treatment data, and iii) both no-treatment and treatment data in Data set 1. Each virtual cohort consists of 10,000 virtual tumors with varying model parameters. For the first virtual cohort, we vary tumor proliferation (*α_m_*), rate of tumor cell death via fast killing (*p*_fast_), and CTL recruitment (*µ*) rates using Latin Hypercube Sampling (LHS) within realistic ranges of parameters. We denote this as the “no data integration (NDI) cohort”. We then construct a second virtual cohort using tumor growth data in mice with no treatment in Data Set 1. We estimate the posterior distributions of *α_m_, p*_fast_ and *µ* by conducting a practical identifiability analysis using the Metropolis-Hastings algorithm. We sample from the posterior distributions instead of the uniform distributions used to generate the NDI cohort. We denote this as the “partial data integration (PDI) cohort”. We create the third virtual cohort using tumor growth data in mice with and without anti-PD-L1 treatment. We denote this as the “complete data integration (CDI) cohort”. In Section 3, we compare the conclusions obtained from these three virtual cohorts, and then use the CDI cohort to test the effects of alternative dosing strategies of anti-PD-L1.

We use Data set 2 to assess the effects of mouse immunocompetency and types of measurement on model identifiability. Data set 2 contains tumor volume measurements of immunocompromised mice and immunocompetent mice at various time points, as well as endpoint CTL density measurement for immunocompetent mice. We use different combinations of these data types to evaluate the the practical identifiability of *α_m_, µ* and *p*_fast_ in a no-treatment model. We construct virtual cohorts using i) only data of immunocompromised mice, ii) tumor volume data of immunocompromised and immunocompetent mice, iii) tumor volume and CTL density data of immunocompetent mice, and iv) all types of data in Data set 2.

We use Data Set 3 to emphasize the importance of modeling the differentiated immune responses to high and low antigen tumor cells. Data Set 3 differs from the first two in that separate *in vivo* experiments were conducted using SIY 1x and SIY 2x tumor cells. Due to the different levels of SIY expressions, we regard SIY 1x tumor cells as LA tumor cells and SIY 2x tumor cells as HA tumor cells. We conduct practical identifiability analysis of *α_m_, µ* and *p*_fast_ in a no-treatment model and create HA and LA virtual cohorts respectively. We then implement the dosing schedule used in our *in vivo* experiments in Data set 1 and evaluate the treatment effects in the HA and LA cohorts.

## 3 Results

### 3.1 Impact of data integration on predictions of therapeutic outcomes

Using Date Set 1, which comprises both no-treatment (control) and treatment *in vivo* data, we construct virtual cohorts using different types of data to simulate one control and two treatment scenarios. We demonstrate the impact of data integration on predictions of therapeutic outcomes and examine the differences between a complete checkpoint blockade and various realistic dosing regimens of anti-PD-L1.

#### Results of practical identifiability analysis

Figure 1A shows model predictions of the temporal changes of tumor volume in the “partial data integration” (PDI) cohort with an active immune checkpoint, plotted in conjunction with no-treatment *in vivo* data. The red lines show the mean tumor volume and standard deviation at each time point of measurement in the pretreatment data. The tumor growth patterns in the virtual cohort closely align with the experimental data. Figure 1B shows the posterior distributions and pairwise heatmaps of *α_m_, p*_fast_ and *µ* following practical identifiability analysis. The heatmap visualizes the posterior joint probability density distribution of *p*_fast_ and *µ*, with brighter regions corresponding to more likely regions for the pairs of parameters. The distribution of *α_m_* in Figure 1B is unimodal, and thus *α_m_* is practically identifiable from no-treatment data. On the other hand, the distributions of *µ* and *p*_fast_ lack a single peak, meaning they are not practically identifiable. The heatmap of the pairwise frequencies of the parameters does not reveal any identifiable combinations of parameters. Nonetheless, we observe the cone shape of the high-frequency regions in the pairwise heatmaps of *α_m_, p*_fast_ and *α_m_, µ*, with the highest frequencies near the mode of *α_m_*. This means that *p*_fast_ and *µ* are likely correlated with *α_m_*. Therefore, if we can fix the value of *α_m_*, the practical unidentifiability of *p*_fast_ and *µ* can be resolved.

**Figure 1:**
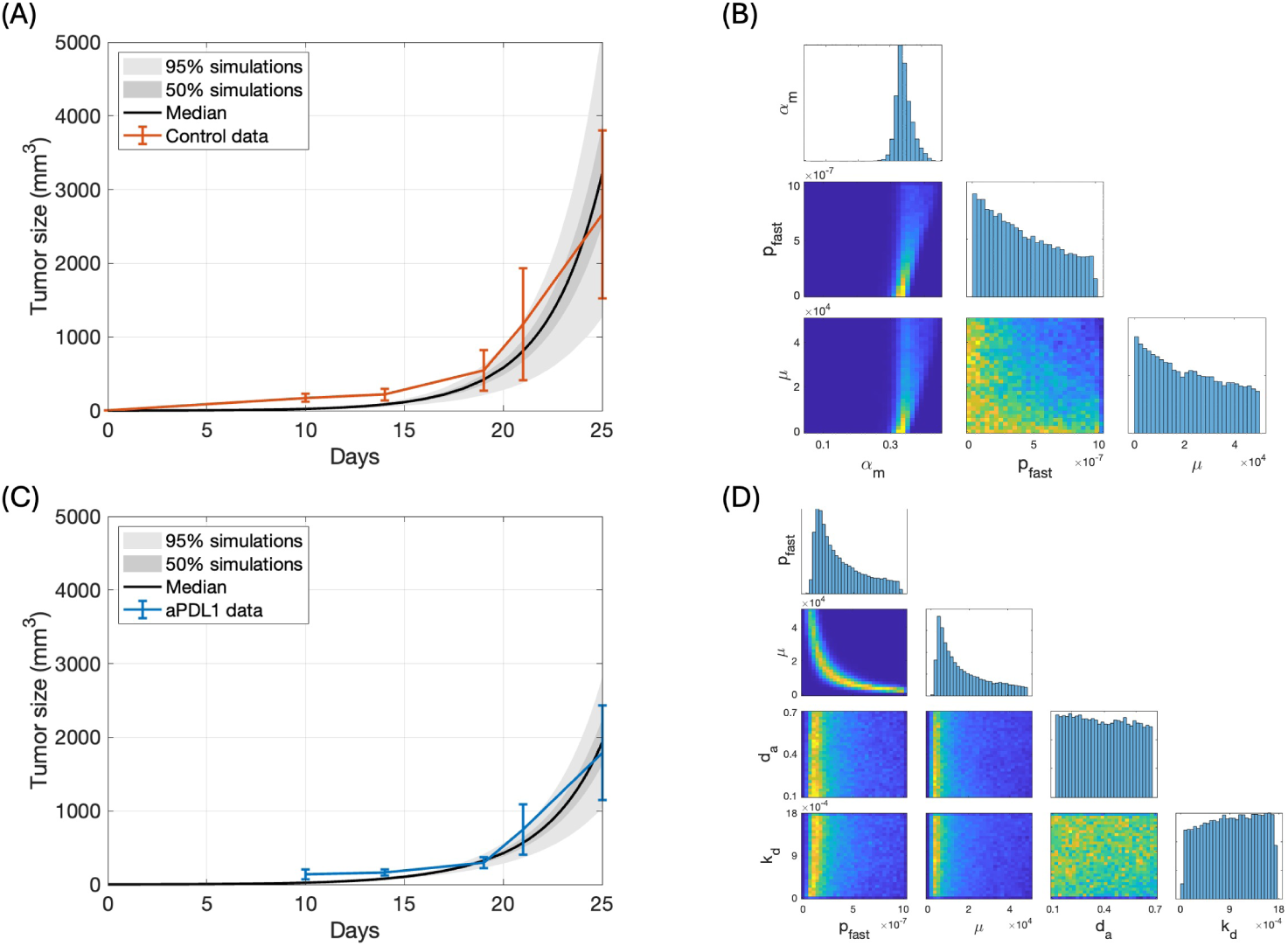
Practical identifiability analysis and virtual cohort construction using Data Set 1. (A) Simulated tumor growths in the PDI cohort with an active immune checkpoint (no treatment) versus no-treatment experimental data. (B) Posterior distribution of model parameters based on no-treatment data, used to sample parameters for constructing the PDI virtual cohort. (C) Simulated tumor growths in the CDI cohort with anti-PD-L1 treatments following our experimental dosing schedule versus treatment experimental data. (D) Posterior distribution of model parameters based on both no-treatment and treatment data, used to sample parameters to construct the CDI virtual cohort. Tumor proliferation rate *α_m_* is fixed at the modal value in (B).

The “complete data integration” (CDI) cohort is constructed using both control and treatment *in vivo* data. The identifiability analysis in Figure 1B shows that *α_m_* is identifiable. We expect tumor proliferation rates to be similar in the no-treatment data and treatment data. Hence, we fix *α_m_* at its unique mode (0.336 per day) to estimate immune parameters *µ* and *p*_fast_, along with treatment-related parameters: natural degradation date of anti-PD-L1 (*d_a_*) and PD-L1-anti-PD-L1 dissociation constant (*k_d_*). Figure 1C shows model-simulated temporal changes of tumor volume in the CDI cohort, plotted alongside experimental measurements of tumor volumes in mice that received anti-PD-L1 treatment following a clinically relevant dosing regimen. The blue lines show the mean tumor volume and standard deviation at each time point of measurement. The simulated post-treatment tumor growths align closely with experimental data. Figure 1D shows the posterior distributions and pairwise heatmaps of the key parameters resulting from the practical identifiability analysis. Unlike the identifiability results using partial data integration shown in Figure 1B, immune parameters *µ* and *p*_fast_ have become identifiable as a pair and demonstrate an inversely proportional relationship. Likely due to improved practical identifiability of key parameters, the range of simulated tumor growth in the CDI cohort is more constrained than in the PDI cohort. The treatment-related parameters *d_a_* and *k_d_*are not practically identifiable, as evidenced by the relatively uniform posterior distribution in the given range.

#### Prediction of percentage tumor reduction

Using the NDI, PDI and CDI virtual cohorts, we create virtual clones to predict the reduction of tumor volume and show that increasing data integration produces more accurate and reliable predictions. Virtual clones are simulated using the ODE model with the same tumor and immune parameters while varying treatment conditions. We simulate tumor growths under one control scenario (active immune checkpoint) and two treatment scenarios: anti-PD-L1 antibody treatment following our experimental dosing schedule resulting in partial checkpoint blockade (directly comparable to post-treatment data), and complete checkpoint blockade (the best possible treatment outcome). We then calculate the reduction in tumor volume due to therapy on day 25 in the two treatment scenarios, compared to the virtual clone in the no-treatment group.

In Figure 2A, green box plots correspond to the percentage change in tumor volume in each virtual cohort after complete checkpoint blockade, and the blue box plots correspond to the results after anti-PD-L1 treatment using the experimental dosing regimen. The first column of Figure 2A corresponds to the simulated therapeutic outcomes of the NDI cohort constructed without experimental data. This cohort predicts a wide range of percentage tumor reduction for both complete checkpoint blockade and anti-PD-L1 treatment. The interquartile ranges of percentage tumor reduction are 39% to 97% after complete checkpoint blockade and 27% to 93% after anti-PD-L1 treatment. The second column of Figure 2A corresponds to the simulated therapeutic outcomes of the PDI cohort constructed using no-treatment data. This cohort predicts similar interquartile ranges of percentage reduction in tumor volume as the NDI cohort, but the median reduction after complete checkpoint blockade is now lower at 72% compared to 83% in the NDI cohort. The third column of Figure 2A corresponds to the CDI cohort constructed using both no-treatment and treatment data. Our model now predicts a significantly narrower interquartile range of tumor reduction under either treatment scenario compared to the NDI and PDI cohorts. The simulated median percentage tumor volume reduction is 55% after a complete checkpoint blockade and 38% after anti-PD-L1 treatment. The interquartile ranges for tumor reduction are 46% to 65% after complete checkpoint blockade and 32% to 45% after realistic anti-PD-L1 treatments. The third cohort predicts no cases of tumor elimination after either treatment.

**Figure 2:**
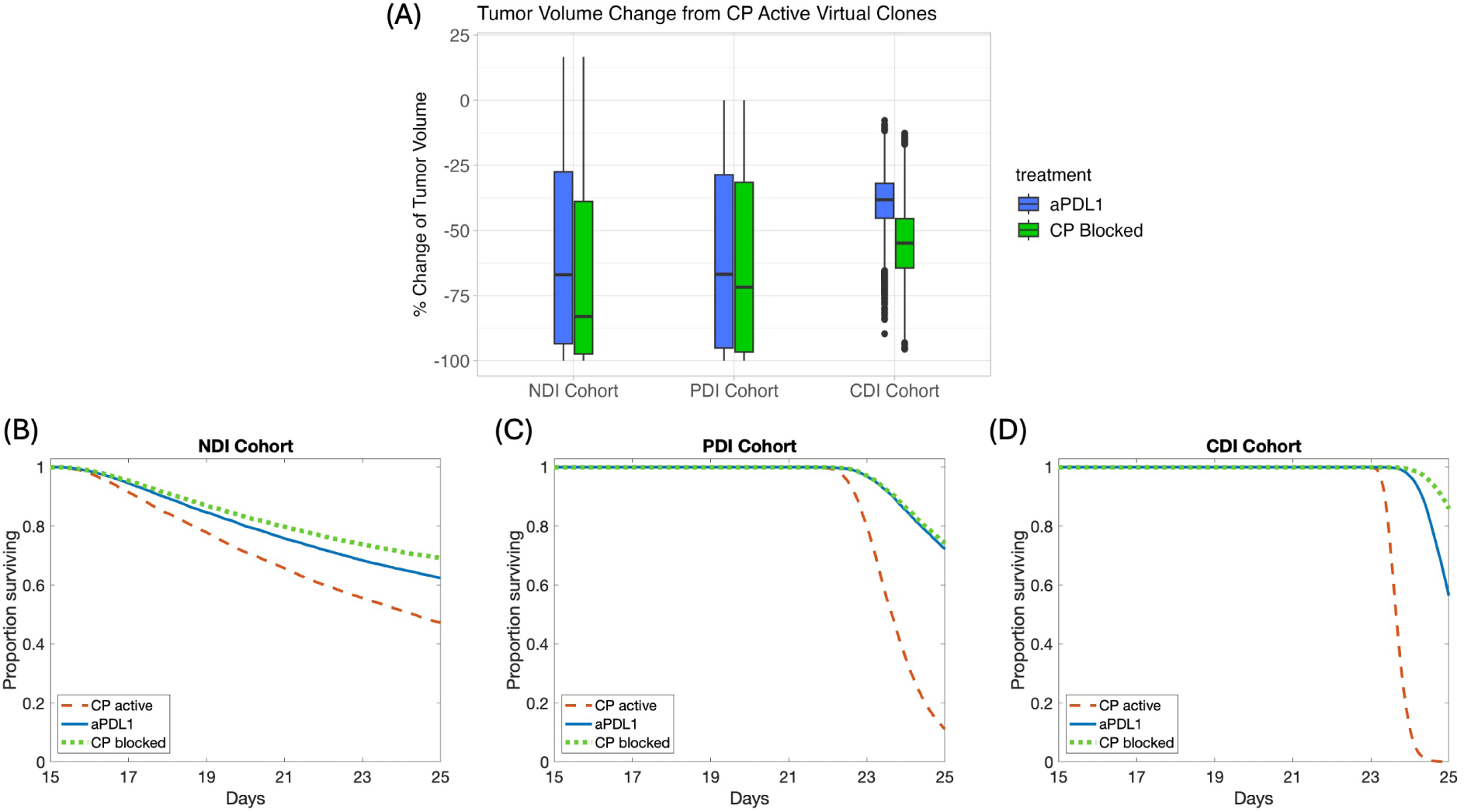
Comparison of model predictions by three virtual cohorts constructed with different extents of data integration, under one control and two treatment scenarios. “CP active” simulates the no-treatment scenario. “APDL1” simulates a partially blocked immune checkpoint under the clinically relevant dosing schedule of anti-PD-L1. “CP blocked” simulates a completely blocked immune checkpoint as the best-case scenario of ICI treatments. (A) Percentages of tumor reduction in two treatment scenarios compared to control virtual clones with an active immune checkpoint. (B) Proportion surviving in the NDI cohort under three scenarios: checkpoint active (red dotted), experimental dosing regimen of anti-PD-L1 (blue solid), and complete checkpoint blockade (green dotted). Death of a virtual mouse is defined as when the tumor volume first exceeds 2000mm^3^; (C) Proportion surviving in the PDI cohort; (D) Proportion surviving the CDI cohort

Note that the blue box plot in the third column most closely reflects tumor reduction *in vivo* because the CDI cohort was constructed such that the simulated pre- and post-treatment tumor growths match the experimental data with and without anti-PD-L1 treatment, respectively. Contrary to predictions by the NDI and PDI cohorts, the CDI cohort predicts no tumor elimination after anti-PD-L1 treatment *in vivo*. When no data or only pretreatment data is available, the first two virtual cohorts falsely predict a nontrivial portion of tumors to be eliminated under either treatment scenario. The virtual NDI and PDI cohorts also overestimate the efficacy of complete checkpoint blockade or anti-PD-L1 treatment because, overall, they predict a greater percentage reduction in tumor volume compared to the CDI cohort. The PDI and CDI cohorts predict that all tumors will shrink in size after complete checkpoint blockade or anti-PD-L1 treatment, but the NDI cohort predicts that some tumors will increase in size after either treatment. As expected, all three cohorts predict a greater reduction of tumor volume after a complete checkpoint blockade compared to anti-PD-L1 treatment following a realistic dosing schedule. However, the difference between the two treatment scenarios is more profound in the CDI cohort, where the interquartile ranges of predicted tumor reduction do not overlap. In general, the CDI cohort constructed using the most amount and types of *in vivo* data gives tighter and more accurate predictions of tumor reduction after immune checkpoint blockade therapy.

#### Prediction of endpoint survival

Data availability and integration into model analysis also affect survival predictions after treatment. We define the “death” of a virtual mouse as the first time the tumor exceeds 2000 mm^3^. This choice aligns with the protocol for sacrificing mice in the experimental studies on which our model is based. Proportion surviving is the ratio of live mice to the total number of mice in the virtual cohort. Figure 2B,C,D show the survival plot of each of the three virtual cohorts constructed above under three scenarios: checkpoint active or no treatment (red dashed line), anti-PD-L1 treatment following the actual experimental dosing schedule (blue solid line) and checkpoint completely blocked (green dotted line). Figure 2B corresponds to the NDI cohort, C to the PDI cohort, and D to the CDI cohort. We recap that the CDI cohort in panel D is the most realistic cohort because it is constructed using no-treatment and treatment *in vivo* data.

All cohorts predict that the endpoint (day 25) proportion surviving is lower with an active immune checkpoint than with either scenario of checkpoint blockade therapy. With an active immune checkpoint, both the PDI and CDI virtual cohorts predict that most or all mice will succumb to cancer by day 25, with the PDI virtual cohort predicting 11.2% surviving by day 25 in panel C, and the CDI cohort predicting no mice surviving beyond day 25 in panel D. On the other hand, the NDI cohort in panel B predicts that close to half (47.3%) of the mice with an active immune checkpoint survive to day 25. The PDI and CDI cohorts more accurately model the *in vivo* data where mice were sacrificed on day 25.

With anti-PD-L1 treatment following the experimental dosing schedule, the endpoint proportion surviving in the PDI and CDI cohorts improve significantly from pretreatment virtual clones to 72.3% in panel C and 56.5% in panel D, respectively. In panel B with the NDI cohort , the endpoint proportion surviving after anti-PD-L1 treatment improves marginally to 62.5%, compared to 47.3% in pretreatment virtual clones. Across all three cohorts, a complete checkpoint blockade improves the endpoint proportion surviving even further, compared to outcomes after anti-PD-L1 treatments. This improvement is most significant in the CDI cohort, with nearly a 30% increase in the proportion surviving, and the smallest in the PDI cohort, with only a 2% increase.

The NDI cohort the least accurate in predicting survival because it dramatically overestimates the endpoint proportion surviving and estimates the first death to occur much earlier than the PDI and CDI cohorts. The NDI cohort predicts the first death to be on day 15, whereas the PDI and CDI cohorts expect the first death to happen between day 21 and day 23. The PDI and CDI cohorts predict the time of first death more accurately because no mouse in the experiments had tumor volume greater than or even close to 2000 mm^3^ on day 15.

Considering that the CDI cohort constructed using both no-treatment and treatment data aligns more closely with reality, the NDI and PDI cohorts overestimate the endpoint survival of the cohorts with an active immune checkpoint or after receiving anti-PD-L1 treatments. The NDI cohort has a much larger error for no-treatment endpoint survival than the PDI cohort. Furthermore, the NDI and PDI cohorts underestimate the difference in the effect on survival of complete checkpoint blockade and anti-PD-L1 treatment following a clinically relevant dose schedule.

### 3.2 Immune parameters and dosing schedules determine post-treatment survival

In this section, we investigate how tumor immune characteristics and alternative dosing schedules influence endpoint (day 25) survival of the virtual cohort. In the most realistic virtual cohort, the complete data integration (CDI) cohort in Figure 2D, we observe that not all mice survive to day 25 even after complete checkpoint blockade, the best-case scenario of ICI treatment. Furthermore, the survival rate after complete checkpoint blockade is approximately 30% higher than after anti-PD-L1 treatment following a realistic dose regimen. All mice that survive with anti-PD-L1 treatment also survive with complete checkpoint blockade treatment.

To investigate the common characteristics of the mice in each post-treatment scenario, we divide the mice in the CDI cohort into three groups: death regardless of treatment (Group R, plotted in red in Figure 3), survival only with complete checkpoint blockade (Group B, plotted in blue), and survival with either anti-PD-L1 treatment or complete checkpoint blockade (Group G, plotted in green). Figure 3A shows the distribution of the CTL recruitment rate *µ* and the fast killing rate *p*_fast_ colored by group. As we discussed in Section 3.1, they are inversely proportional and approximately follow the relationship *µ* · *p*_fast_ = *k*. We denote the best-fit proportionality constants for each group as *k_r_, k_b_, k_g_*, which satisfy *k_r_ < k_b_ < k_g_* from the figure. Therefore, when the product *µ* · *p*_fast_ is small, as is the case in Group R, the mouse is likely to die by day 25 despite receiving checkpoint blockade therapy in the best-case scenario. When *µ* · *p*_fast_ is large, Group G, the mouse will likely survive after receiving anti-PD-L1 treatment following our experimental dosing schedule. For values of *µ* · *p*_fast_ in between (Group B), the mouse survives if the checkpoint is completely blocked but dies with our current anti-PD-L1 dosing regimen.

**Figure 3:**
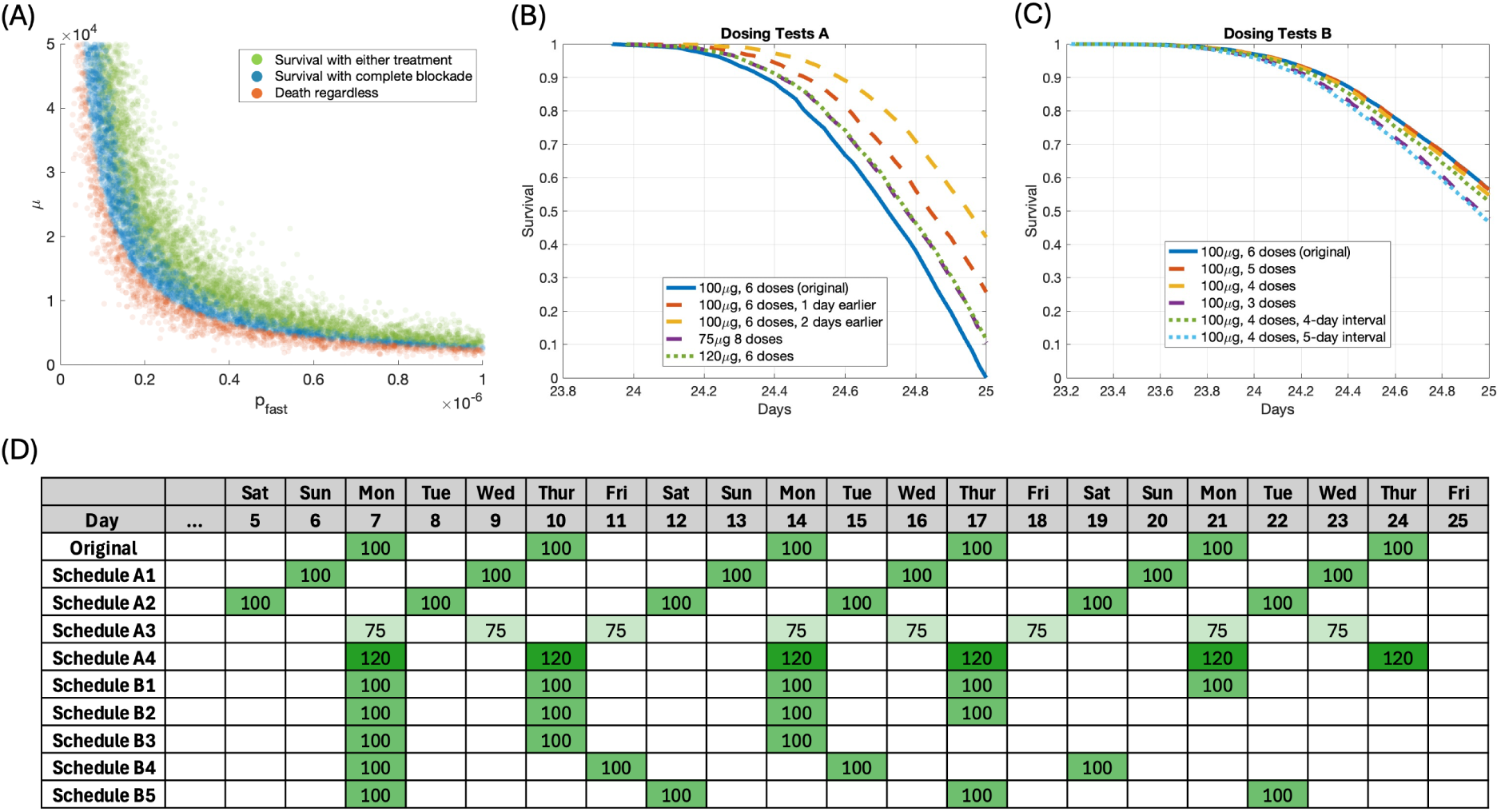
Therapeutic outcomes in the most realistic CDI cohort under various anti-PD-L1 treatment scenarios (A) Distribution of immune parameters stratified by treatment outcomes. (B) Alternative dosing schedules, set A, on Group B, which survive to day 25 following complete checkpoint blockade but not under our experimental anti-PD-L1 dosing schedule. Schedules A1-A3 deliver the same total dose as the original schedule, but vary in treatment start time. Schedule A4 matches the original timing but administers 1.2 times the total dose. (C) Alternative dosing schedules, set B, on 10 000 randomly selected samples in the CDI cohort. All schedules have the same treatment start time as the original schedule but differ in the number of doses and/or intervals between doses. (D) Original and alternative dosing schedules.

We first focus on Group B to test whether alternative dosing schedules can improve the survival proportion on day 25. In our *in vivo* treatment study, mice received 100 µg of anti-PD-L1 antibody on Days 7, 10, 14, 17, 21, and 24, for a total cumulative dose of 600 µg. Despite this treatment regimen, no long-term survivors were observed. On the other hand, complete checkpoint blockade would result in a 100% survival rate in Group B. We vary dosing schedules along two primary dimensions: dosage and timing. Since biological efficacy of checkpoint blockade saturates after the maximum dose, our computational tests focus primarily on modifying the timing of administration and evaluating the effects of lower doses. Alternative schedule A1 (red, dashed line) and A2 (yellow, dashed line) shift the original schedule (blue, solid line) one day or two days early, respectively, and keep the dosage of each administration the same. Alternative schedule A3 (purple, dashed line) starts the first dose on the same day as the original schedule (day 7), and administers smaller and more frequent doses. Each dose is reduced to 75 µg, compared to the original 100 µg, while maintaining the same total cumulative dosage as the original schedule. As Figure 3B shows, shifting the schedule by one day improves the endpoint survival of Group B from 0% to 26%. Shifting the schedule by two days further improves the survival by 16%. Smaller and more frequent doses that start on day 7 improves the survival of Group B from 0% to 10% while keeping the total dosage the same. Alternative schedule A4 (green, dotted line), which follows the same timing as the original regimen but increases the dosage by 1.2-fold, is not clinically feasible and is included for comparison purposes only. The results indicate that a higher dosage does not necessarily improve endpoint survival; in fact, when compared to schedules A1 and A2, schedule A4 yields inferior outcomes.

We then expand our analysis to the entire virtual cohort. In addition to maximizing endpoint survival, we also aim to identify strategies that reduce the total dosage administered without compromising overall survival outcomes. We evaluate the effects of shortening the original six-dose schedule by stopping treatment after five (Schedule B1), four (Schedule B2), and three (Schedule B3) doses, respectively. The survival curve is plotted in Figure 3C, with the blue solid line corresponding to the original schedule, red, yellow and purple dashed lines corresponding to Schedules B1, B2 and B3 respectively. The decrease in survival from six to five doses is negligible, and the decrease from five to four doses is 1.7%. In comparison, the drop in survival from four to three doses is significantly larger at 7%. Using a total of four doses, we implement Schedules B4 and B5 to examine the impact of extended intervals between administrations. In the original schedule, doses are administered every three days, with slight adjustments for weekends. In contrast, Schedule B4 uses a four-day interval between doses, while Schedule B5 extends the interval to five days. Schedule B4 results in approximately a 4% reduction in survival compared to the baseline, and Schedule B5 leads to an additional 6% decrease. Notably, Schedule B5 (4 doses extended interval) and Schedule B3 (3 doses, original interval) have nearly indistinguishable survival curves.

### 3.3 Additional experiments enhance practical identifiability

Using Data Set 1, we showed that the complete data integration (CDI) cohort yields the most reliable predictions of ICI treatment outcomes. The key distinction between the CDI cohort and the partially (PDI) or no-data integration (NDI) cohorts lies in the identifiability of the CTL recruitment rate (*µ*) and the fast-killing tumor cell death rate (*p*_fast_). We utilize Data Set 2 — which includes experiments involving immunocompromised and immunocompetent mice, tumor volume measurements, and endpoint CTL density — to demonstrate how incorporating diverse types of experimental data, beyond just treatment and control comparisons, can enhance model identifiability and accurately predict therapeutic outcomes of ICIs.

We first performed a practical identifiability analysis as a control using only tumor volume measurements in immunocompetent mice. Figure 4A shows the estimated posterior distributions. Similar to those in Figure 1B, the tumor proliferation rate *α_m_*is identifiable. To evaluate how the additional experiments and measurements enhance the identifiability of the model, we conduct three additional rounds of analysis, adding RAG1 KO data and endpoint CTL measurements as observables one at a time and then combined. We first used tumor volume measurements from

**Figure 4:**
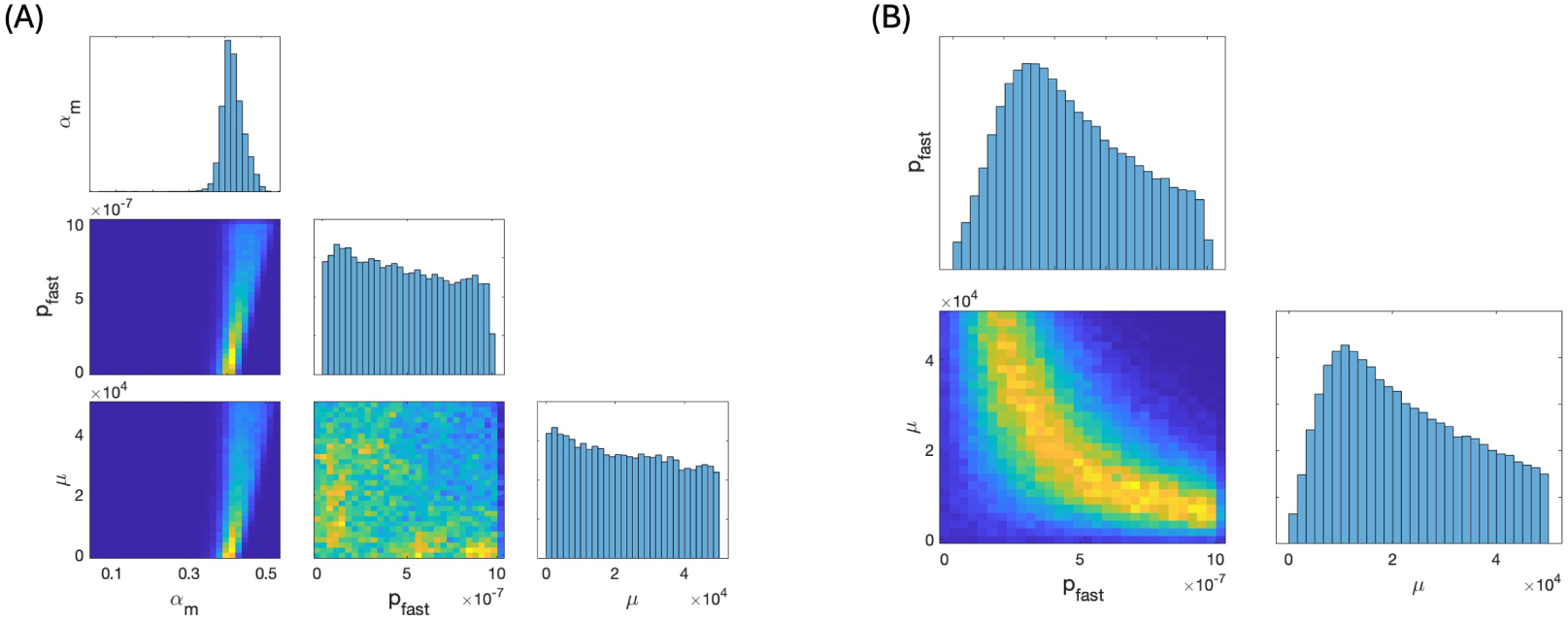
Practical identifiability analysis of the ODE model using different types of data in Data Set 2. (A)Use tumor volume measurements in C57BL/6J mice. Only tumor proliferation rate (*α_m_*) is identifiable. (B) Use tumor volume measurements in RAG1 KO mice to estimate *α_m_* and use tumor volume measurements in C57BL/6J mice to estimate immune parameters. Immune parameters *µ* and *p*_fast_ are identifiable as a pair.

RAG1 KO mice to estimate the distribution of tumor proliferation rate using a simple exponential growth model (equation S3). Figure S2A shows the estimated posterior distribution. The tumor proliferation rate *α_m_*is identifiable with a single mode of 0.424 d^−1^. Figure S2B shows the simulated tumor growths and the RAG1 KO experimental data. Since the same type of tumor cells is used for immunocompetent mice, we assume that tumor cells have the same proliferation rate and fix *α_m_*at 0.424 d^−1^. We then use the MCMC Metropolis-Hastings algorithm to estimate the distribution of *p*_fast_ and *µ*. Similar to Figure 1D for Data Set 1, the tumor cell death rate by fast killing (*p*_fast_) now exhibits a single mode centered around 3.13 × 10^−7^, and the CTL recruitment rate (*µ*) has a single mode near 10,625. Pairs of *p*_fast_ and *µ* are now identifiable. As illustrated in the heatmap in Figure 4B, the two parameters appear inversely correlated. We notice that simulated tumor growths in the virtual cohort created by sampling the distributions in Figure 4A is almost indistinguishable from the cohort created by sampling the distributions in Figure 4B, as shown in Figure S3. This suggests that virtual cohorts generated using either practically identifiable or unidentifiable immune parameters can produce no-treatment tumor growth trajectories that fit the experimental data equally well and appear nearly indistinguishable.

To assess whether endpoint CTL measurements in immunocompetent mice can similarly enhance the practical identifiability of immune parameters, we combine CTL density and tumor volume measurements from C57BL/6J mice to estimate the distributions of *α_m_, µ* and *p*_fast_. The results, shown in Figure S4A, are similar to Figure 4A. Only *α_m_* is practically identifiable. With both endpoint CTL measurements and RAG1 KO data, we can also fix *α_m_* to estimate the distributions of *p*_fast_ and *µ*, as we do in Figure 4B. The results of the practical identifiability analysis shown in Figure S4A is also similar to Figure 4B. The *µ* − *p*_fast_ heatmap does not reveal a more constrained or well-defined region of high likelihood. Thus, in this case, the addition of endpoint CTL measurements has minimal impact on improving the identifiability of the immune parameters. Estimating *α_m_* with confidence from the RAG1 KO data and fixing it to the single-modal value is crucial for the practical identifiability of *µ* − *p*_fast_.

### 3.4 Tumor antigenicity and associated immune characteristics determine responses to ICIs

Through practical identifiability analysis using Data Set 3, we construct high antigen (HA) and low antigen (LA) virtual cohorts and implement the clinically relevant, experimental dosing schedule of anti-PD-L1. Results suggest that the differing responses to ICI treatments between the two cohorts are driven primarily by distinct immune characteristics, rather than variations in tumor proliferation rates — highlighting the critical role of immune context in shaping ICI treatment outcomes.

#### Construction of HA and LA virtual cohorts

Using SIY 1x and SIY 2x *in vivo* data, we perform a practical identifiability analysis and generate HA and LA virtual cohorts as described in Section 2.5. The cohorts feature an active immune checkpoint and accurately reproduce experimental observations under no-treatment conditions, as shown in Figure 5. Despite a wide range of key immune parameters (*p*_fast_ and *µ*) used to generate the virtual cohorts, the simulated growths with an active immune checkpoint show a narrow 95% simulated interval and closely align with the experimental data. Results of the practical identifiability analysis is similar to only using pretreatment data in Data Set 1 (Figure 1B) and only using tumor volume of immunocompetent mice in Data Set 2 (Figure 4A): only *α_m_* is identifiable, as shown in the posterior distributions of parameters in Figure 5B and D. The median tumor proliferation rate (*α_m_*) for HA tumors (0.274 d^−1^) is smaller than that for LA tumors (0.288 d^−1^), agreeing with our observation that HA tumors grow slower than LA tumors in Figures 5A and C.

**Figure 5:**
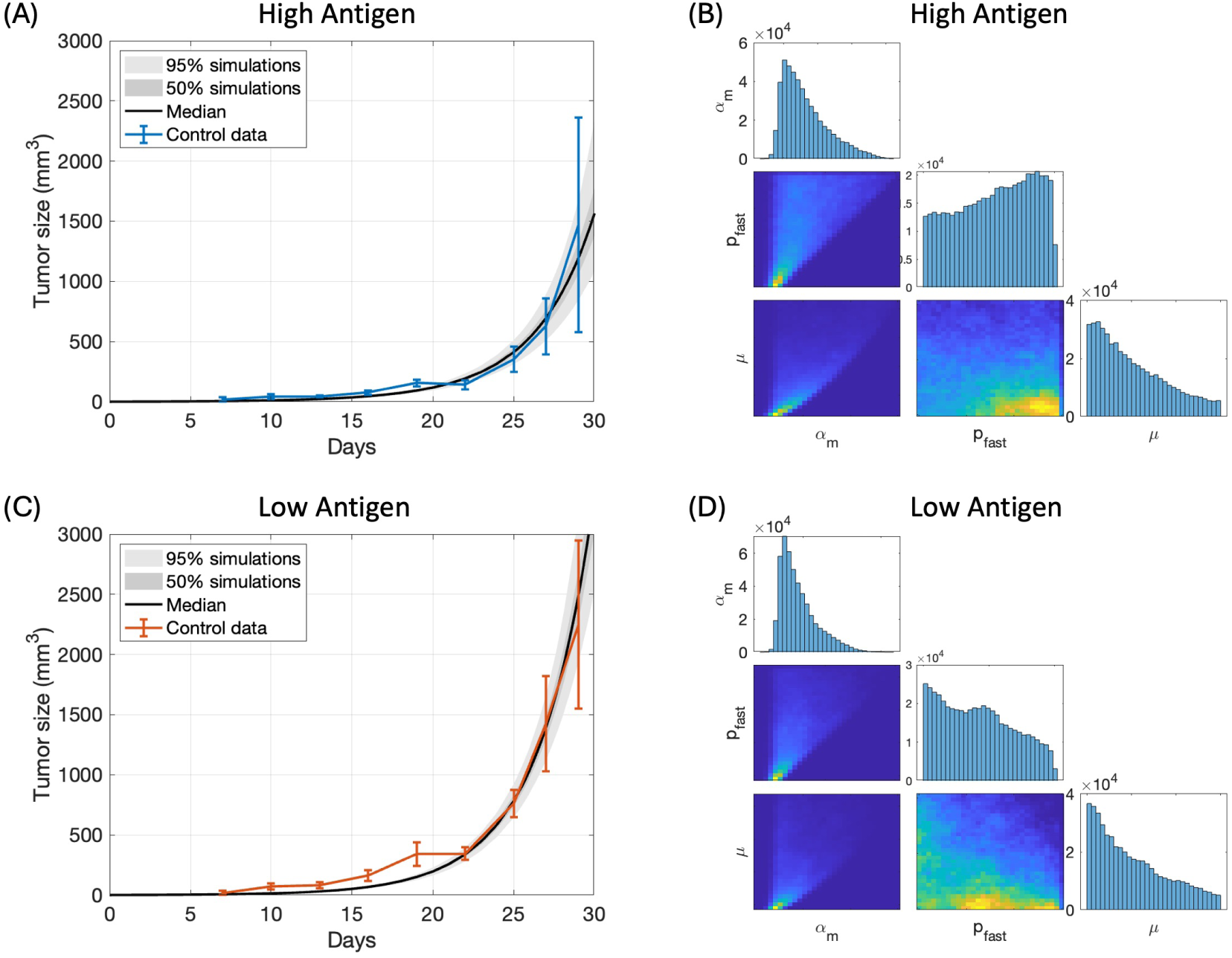
Practical identifiability analysis and virtual cohorts using Data Set 3. (A) Simulated SIY 2x (HA) tumor growths with an active immune checkpoint vs pretreatment experimental data. (B) Posterior distribution of parameters of HA tumors. (C) Simulated SIY 1x (LA) tumor growths with an active immune checkpoint vs pretreatment experimental data. (D) Posterior distribution of parameters of LA tumors.

#### Tumor antigenicity affects efficacy of anti-PD-L1 treatment

We then compare the efficacy of anti-PD-L1 treatment for the HA and LA virtual cohorts by implementing the dosing schedule used in our *in vivo* experiments in Data Set 1. Following anti-PD-L1 treatment, all tumors in both virtual cohorts exhibit regression. Although pretreatment tumor growth is nearly identical across cohorts, as shown in Figures 5A and C, the response to immune checkpoint inhibition varies considerably within each cohort. This is reflected in the wide distribution of endpoint tumor volumes after treatment, as illustrated in Figure 6A. The HA cohort displays a narrower range of post-treatment tumor sizes, with tumors generally smaller than those in the LA cohort, suggesting greater overall sensitivity to ICI therapy. The median endpoint tumor size in the HA cohort (74 mm^3^) is significantly smaller than that in the LA cohort (418 mm^3^). Given that pretreatment, endpoint HA tumors are smaller than LA tumors, we also evaluate the percentage change in endpoint tumor volume relative to their corresponding virtual clones with an active immune checkpoint, as shown in Figure 6B. The percentage reduction of tumor size in the HA cohort is greater than in the LA cohort, with a median of 94% in the HA cohort and a median of 83% in the LA cohort. More than 67% of tumors in the HA cohort achieve a reduction of more than 75% while only 55% of the tumors in the LA cohort achieve the same level of reduction. This shows that anti-PD-L1 treatment is more effective for tumors of high antigenicity than tumors of low antigenicity.

**Figure 6:**
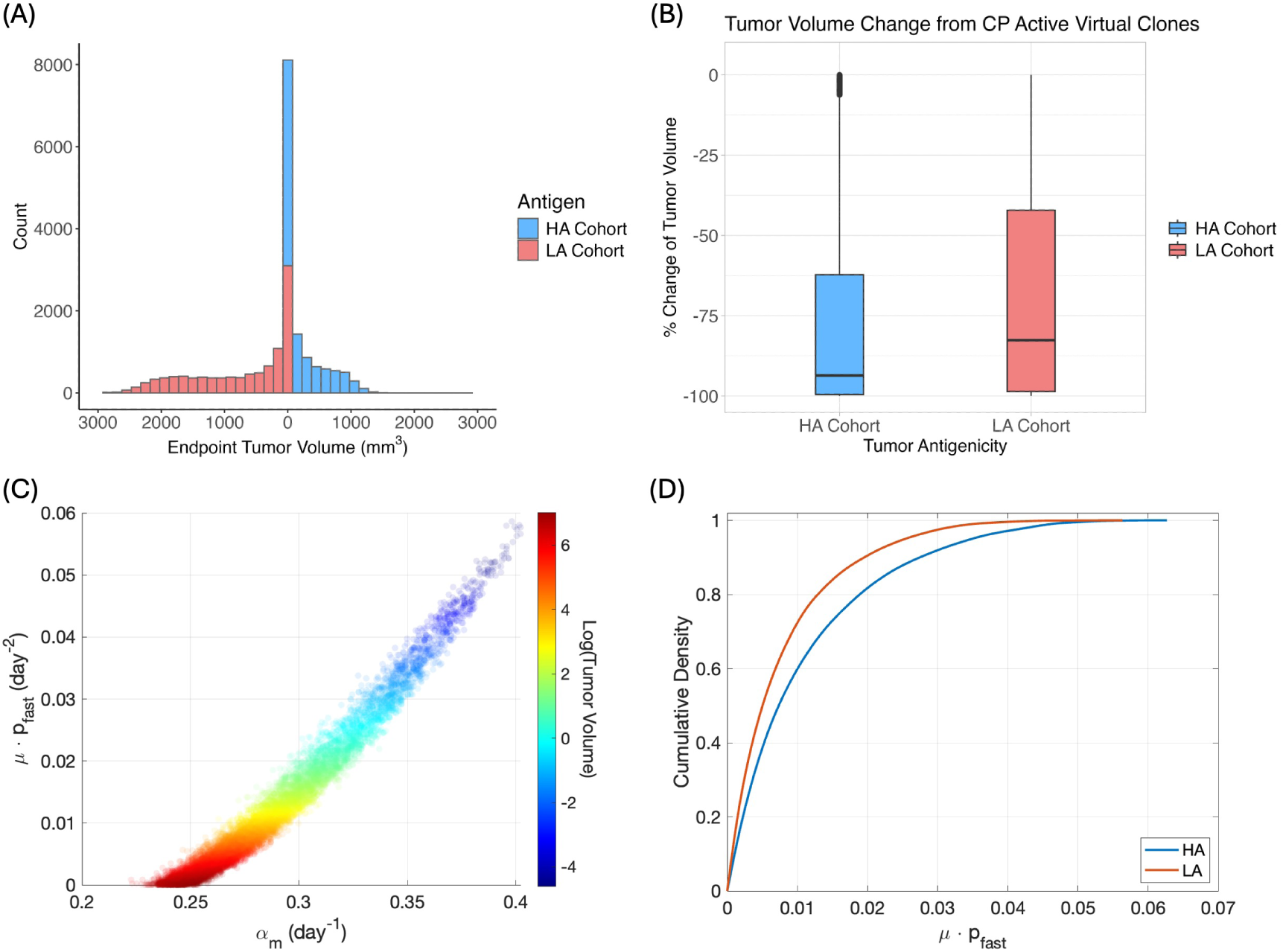
Differed post-treatment tumor outcomes of HA and LA virtual cohorts and potential reasons for the differentiated immune responses. (A) Distribution of endpoint (Day 29) tumor volumes in the HA and LA virtual cohorts, respectively, after anti-PD-L1 treatment following our experimental dosing schedule. (B) Percentage tumor volume change in the post-treatment virtual cohorts compared to their pretreatment virtual clones. (C) Affine relationship between *µ* · *p*_fast_ and *α_m_* in the HA cohort, as well as the relationship between the parameters and endpoint tumor volumes after treatment. The color of the dots represents the log of the tumor volume, with red being the largest tumors and purple being the smallest tumors. (D) Cumulative distribution function (CDF) of *µ* · *p*_fast_ in the HA and LA cohorts.

To understand what characteristics of HA and LA tumors lead to disparate responses to anti-PD-L1 treatment, we investigate the relationship between model parameters and the endpoint tumor size. One parameter related to the tumor cell growth (*α_m_*) and two parameters related to the immune system (*µ, p*_fast_) are varied in the virtual cohorts. Figures 1D and 4D suggest that, given a set of experimental data, *µ* and *p*_fast_ likely exhibit an inversely proportional relationship. Therefore, in Figure 6C, we plot the product of the two immune parameters *µ* · *p*_fast_ against *α_m_* to compare the impact of immune parameters with that of tumor proliferation parameters. The color of the dots on the scatter plot represents the day 29 tumor size on a log scale after anti-PD-L1 treatment. We include only the HA cohort plot, as the LA cohort’s corresponding plot appears qualitatively similar. We observe that the smallest tumors (in purple) have larger *µ* · *p*_fast_ and larger *α_m_* than the biggest tumors (in red). It may seem counterintuitive that tumors with lower proliferation rates (*α_m_*) exhibit greater growth following ICI treatment. While it is true that, all else being equal, higher proliferation rates lead to larger tumors, the practical identifiability analysis reveals a trade-off. To ensure that simulated tumor growth aligns well with experimental data, compensatory relationships emerge between *α_m_* and immune-related parameters such as *µ* or *p*_fast_. As a result, slower-growing tumors may be paired with weaker immune responses, ultimately leading to larger post-treatment tumor sizes. In fact, as Figure 6C shows, there is a clear positive, almost affine correlation between *α_m_* and *µ* · *p*_fast_. This means that given what we observe experimentally, a mouse cannot simultaneously have a low tumor proliferation rate, high immune recruitment rate and high rate of fast killing. The color bands appear horizontal, implying that the effect of immune parameters (*µ*·*p*_fast_) on the outcomes of checkpoint blockade therapy dominates the impact of tumor proliferation rates (*α_m_*). Therefore, although low tumor proliferation rates and tumor escape after ICI treatment are correlated, this relationship is not causal.

Figure 6D shows the estimated cumulative distribution functions (CDF) of *µ* · *p*_fast_ in the HA and LA cohorts. For a given *x*, the y-value of the CDF should be interpreted as the proportion of the population with *µ* · *p*_fast_ *< x*. We can see that for most *x*, the LA cohort (in red) has more mice with *µ* · *p*_fast_ *< x*, meaning that the LA cohort has smaller *µ* · *p*_fast_ in general. Moreover, a given y-value can be seen as the bottom y-th percentile of the population in terms of *µ* · *p*_fast_. The same percentile in the LA cohort has a lower *µ* · *p*_fast_ than the HA cohort. For example, the median of the HA cohort (0.0072) is about 1.5 times the median of the LA cohort (0.0049). Overall, our analysis confirms that the HA cohort’s better responses to anti-PD-L1 treatment are primarily attributable to their higher immune recruitment rate and higher rate of fast killing compared to the LA cohort.

## 4 Discussion

Understanding and predicting the efficacy of immune checkpoint inhibitor (ICI) therapy remains a major challenge in cancer immunology, particularly due to the complexity and variability of tumor-immune dynamics. Reliable prediction of immunotherapy outcomes depends not only on the biological realism of computational models but also on the extent to which key model parameters can be identified from available data. In this study, we present a systematic, data-driven framework that links structural and practical identifiability analyses with experimental design, model calibration, and treatment optimization in the context of ICI therapy for bladder cancer.

Using three distinct *in vivo* datasets, we show how different levels of data integration — ranging from no data to control-only to full treatment datasets — affect the ability to estimate immune parameters and accurately forecast treatment outcomes. Our findings underscore a central theme: practical identifiability of key immune parameters is essential for making trustworthy quantitative predictions about tumor reduction, survival, and the efficacy of alternative dosing schedules.

Our workflow begins with global sensitivity analysis and structural identifiability analysis, both of which can be conducted before any experimental data are collected. Sensitivity analysis identifies which parameters most strongly influence model outputs, thereby guiding experimental focus and model simplification. Structural identifiability analysis determines whether model parameters are theoretically estimable from ideal data. Because structural identifiability is a prerequisite for practical identifiability, we addressed model ambiguities by combining parameters into effective quantities and fixing low-sensitivity parameters to literature-based values. This stepwise refinement was critical for making the model tractable and for focusing inference on biologically relevant mechanisms.

Our results demonstrate that data integration and practical identifiability are inextricably linked, and that both are required for generating accurate, quantitative predictions of ICI treatment outcomes. Virtual tumor cohorts constructed without treatment data systematically overestimated therapeutic efficacy, predicted unrealistically high tumor clearance rates, and underestimated survival differences between treatment regimens. These failures arose from strong parameter correlations and practical unidentifiability, highlighting that good fits to baseline tumor growth data alone are insufficient to validate predictive models.

When practical identifiability is compromised, collecting additional experimental data may resolve unidentifiability. We demonstrate that immunodeficient control data are more useful than endpoint CTL density measurements for enhancing practical identifiability of key immune parameters in our ODE models. Targeted experimental design is important, although no method guarantees practical identifiability a priori because data quality varies in reality. In case of practical unidentifiability, models can still offer valuable qualitative insights even though their quantitative predictions may not be precise or accurate.

Once practical identifiability is achieved, virtual cohorts become reliable tools for *in silico* optimization. Using cohorts with identifiable parameters, we show that earlier initiation of anti-PD-L1 treatment and smaller, more frequent dosing improve endpoint survival without increasing the total dose. Strategically stopping treatment early preserves overall survival outcomes while lowering the total dose. These results also suggest that immune recruitment and tumor cell-killing capacity define an upper bound on the efficacy of ICI monotherapy. When immune function is weak, schedule optimization alone provides limited benefit, indicating a need for combination strategies that enhance immune activation. A notable limitation of this study is that only one dataset includes both pretreatment and post-treatment data, requiring us to derive and validate practical identifiability within the same dataset. However, these findings align with *in vivo* [16] and clinical [22, 24] observations that lower tumor burden leads to improve efficacy of ICIs. Future work should involve experimental designs that separate calibration from validation—enabling model parameters to be estimated on one dataset and predictions to be validated on another.

Finally, using antigen-high and antigen-low tumor models, we demonstrated that immune parameters more strongly govern differential treatment responses than intrinsic tumor growth rates. This result not only confirms prior modeling studies [56, 55] but does so for the first time using experimental data, highlighting the importance of integrating immune phenotyping into preclinical models of ICI therapy.

## 5 Conclusion

This study provides a comprehensive, data-driven roadmap for improving the predictive reliability of immunotherapy models. By integrating sensitivity analysis, structural identifiability, and practical identifiability with targeted experimental design, we demonstrate how mathematical models can move beyond qualitative insight to quantitative prediction. Across three *in vivo* datasets, we show that sufficient and appropriate data integration is essential for identifying immune parameters that govern treatment response. Models calibrated without identifiable parameters may reproduce pretreatment dynamics yet substantially overestimate therapeutic benefits.

## S6 Supplementary materials

### S6.1 Parameters of ODE models

**Table 1:** Baseline parameters for the ODE model.

| Name | Description | Value (Baseline) | Units | Source |
| --- | --- | --- | --- | --- |
| $\alpha_n$ | Proliferation rate of high antigen tumor cells | 0.05 - 0.8 (0.337) | per day | Estimated ([39]) |
| $\alpha_m$ | Proliferation rate of low antigen tumor cells | 0.05 - 0.6 (0.337) | per day | Estimated ([39]) |
| K | Carrying capacity for tumor cells | $3 - 6 \cdot 10^9 (5 \cdot 10^9)$ | # of cells | Estimated |
| $\delta_{ns}$ | Maximum CTL-induced death rate of high antigen tumor cells via the slow killing mechanism | 1 - 12 (4) | per day | Estimated ([4]) |
| $\delta_{ms}$ | Maximum CTL-induced death rate of low antigen tumor cells via the slow killing mechanism | 1 - 12 (4) | per day | Estimated ([4]) |
| $\delta_{nf}$ | CTL-induced death rate of high antigen tumor cells via the fast-killing mechanism | $10^{-8} - 10^{-6} (2.5 \cdot 10^{-7})$ | per cell per day | Estimated ([39]) |
| $\delta_{mf}$ | CTL-induced death rate of low antigen tumor cells via the fast-killing mechanism | $10^{-8} - 10^{-6} (2.5 \cdot 10^{-7})$ | per cell per day | Estimated ([39]) |
| $\kappa_0$ | Half-saturation constant in maximum CTL-induced death rate of via the slow killing mechanism | $10^6 - 10^8 (2 \cdot 10^7)$ | # of cells | Estimated |
| $\kappa_1$ | Saturation effect by immune cells on slow killing | 0.01 - 1 (0.5) | dimensionless | Estimated ([25]) |
| $\kappa_2$ | Half-saturation constant in the antigen-mediated T cell proliferation rate | $10^6 - 10^8 (2.019 \cdot 10^7)$ | # of cells | Estimated ([39]) |
| $\delta_t$ | Death rate of T cells | 0 - 0.5 (0.0412) | per day | [37] ([25]) |
| $\mu$ | Activation and recruitment rate of T cells | $5 \cdot 10^3 - 1.5 \cdot 10^5 (2 \cdot 10^4)$ | # per day | Estimated |
| $\delta_n$ | CTL death rate due to interactions with high antigen tumor cells | $10^{-11} - 10^{-9} (3.422 \cdot 10^{-10})$ | per cell per day | Estimated ([25]) |
| $\delta_m$ | CTL death rate due to interactions with low antigen tumor cells | $10^{-11} - 10^{-9} (3.422 \cdot 10^{-10})$ | per cell per day | Estimated ([25]) |
| $p_1$ | Probability of high antigen tumor cells death via the fast-killing mechanism | 0 - 1 (0.92) | dimensionless | Estimated |
| $p_2$ | Probability of low antigen tumor cells death via the fast-killing mechanism | 0 - 1 (0.33) | dimensionless | Estimated |
| $\alpha_{nt}$ | Maximum rate of CTL proliferation activated by high antigen tumor cells | 0 - 0.5 (0.15) | per day | Estimated |
| $\alpha_{mt}$ | Maximum rate of CTL proliferation activated by low antigen tumor cells | 0 - 0.5 (0.15) | per day | Estimated |
| $\rho_p$ | Molar concentration of PD-1 per T cell | $10^{-12} - 10^{-10} (1.259 \cdot 10^{-11})$ | $\mu M$ | [49], [37] |
| $\rho_l$ | Molar concentration of PD-L1 per T cell | $10^{-12} - 2 \cdot 10^{-10} (2.510 \cdot 10^{-11})$ | $\mu M$ | [49], [37] |
| $\epsilon_c$ | Expression of PD-L1 on tumor cells vs. T cells | 1 - 50 (10) | dimensionless | [49] |
| $\mu_{PA}$ | Blocking rate of PD-1 by anti-PD-1 | 6.45 - 273 (8.945) | L/$\mu$ mol/h | [49], [28] |
| $k_{TQ}$ | Inhibition of T cells by PD-1/PD-L1 | $10^{-10} - 10^{-8} (1.296 \cdot 10^{-9})$ | $(\mu M)^2$ | [49], [28] |

### S6.2 Identifiability analysis

#### S6.2.1 Structural identifiability

A vector *s*(*p*) is an exhaustive summary of an experiment if it contains only information about the parameters *p* that can be extracted from knowledge of the control *u*(*t*) and the measured quantities *y*(*t*; *p*). The generating series method for structural identifiability assumes that the observables can be expressed as a series expansion with respect to time and inputs, with the series coefficients being *h*(*t, p, x*_0_) and the successive Lie derivatives along the vector fields f and g, evaluated at the initial conditions *x*_0_(*p*). [9] The Lie derivative of h along f is defined as the following:

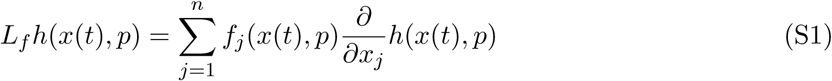

In genSSI 2.0, the exhaustive summary consists of these series coefficients, which form a system of equations in parameters *p*. The structural identifiability of the model depends on whether there is a unique solution to this system of equations. The Jacobian matrix of the exhaustive summary is then calculated with respect to *p*. The Jacobian has as many columns as parameters, and as many rows as non-zero series coefficients. When the rank of the Jacobian matrix is equal to the number of parameters, at least local identifiability is guaranteed. A complete identifiability tableau visualizes the dependencies between parameters by representing the Jacobian matrix as a “0-1” matrix. An element at the (*i*; *j*) position in the identifiability tableau has value “1” if the non-zero series coefficient *L_i_* depends on the parameter *p_j_*, and “0” otherwise. Black squares in an identifiability tableau represent “1” and the white squares represent “0”. In general, the Jacobian has more rows than parameters. A reduced identifiability tableau consists of a number of rows of linearly independent Lie derivatives equal to the generated rank of the extended Jacobian, represented in a similar fashion as the complete identifiability tableau. [9]

**Figure S1:**
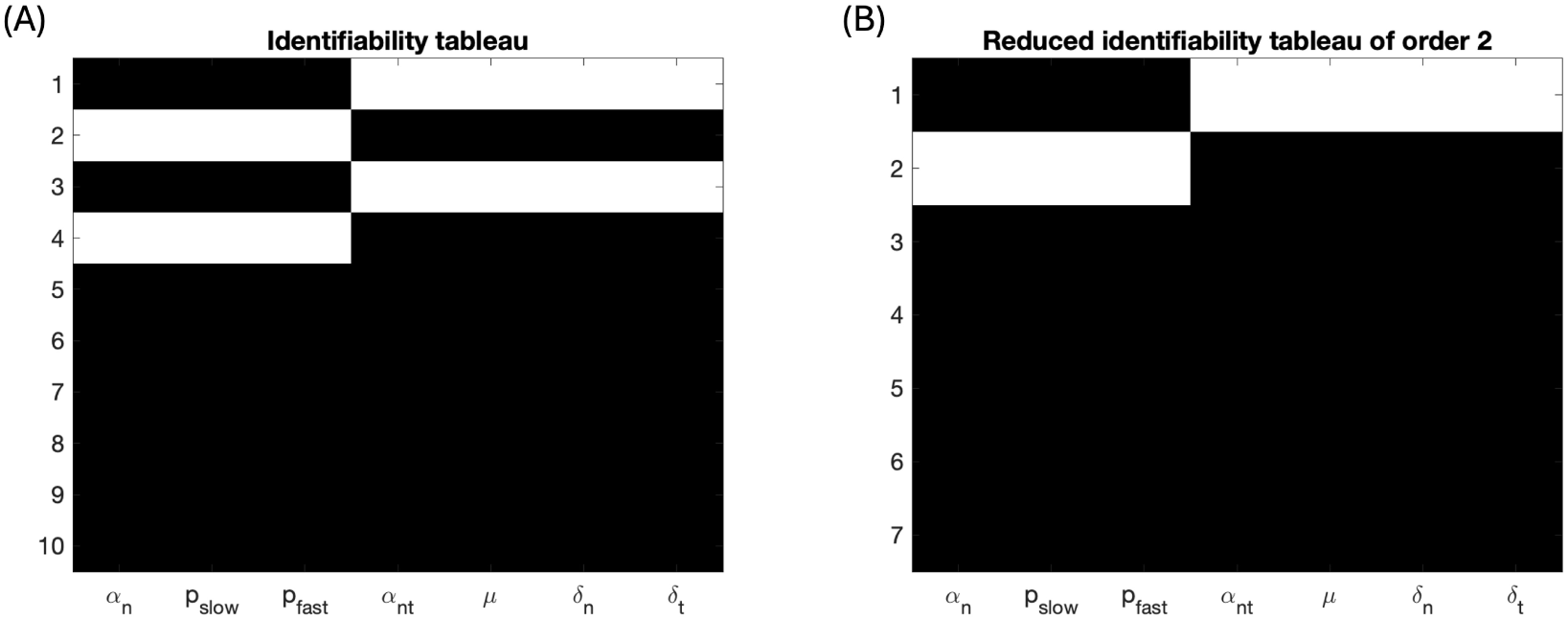
Structural identifiability analysis using genSSI 2.0: identifiability tableaux of the updated, structurally identifiable one-phenotype ODE model with completely blocked or active immune checkpoint

#### S6.2.2 Practical identifiability

At the heart of practical identifiability analysis is estimating the distribution of model parameters given observations. Suppose *p* represents model parameters and *z* represents observed data. By Bayes’ Theorem,

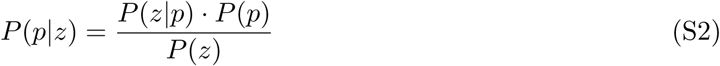

The Metropolis-Hastings algorithm [20, 36] estimates *P* (*p*|*z*), the posterior distribution, by sampling more frequently in regions with higher likelihood (*P* (*z*|*p*)) without having to estimate *P* (*z*). When the Markov Chain converges, the sample distribution approximates the posterior distribution. We implement the Metropolis-Hastings algorithm in MATLAB and analyze the practical identifiability of our ODE models using the *in vivo* data sets. We assume that the prior distributions (*P* (*p*)) are uniform. To ensure that the Markov chain reaches its stationary distribution and accurately approximates the posterior distributions, we initiate the Markov chains from at least four randomly selected locations in the parameter space. Each Markov chain has a minimum length of 3 × 10^4^ and includes a burn-in period of 1 × 10^4^.

### S6.3 Data integration

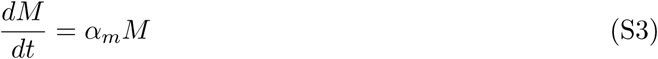

**Figure S2:**
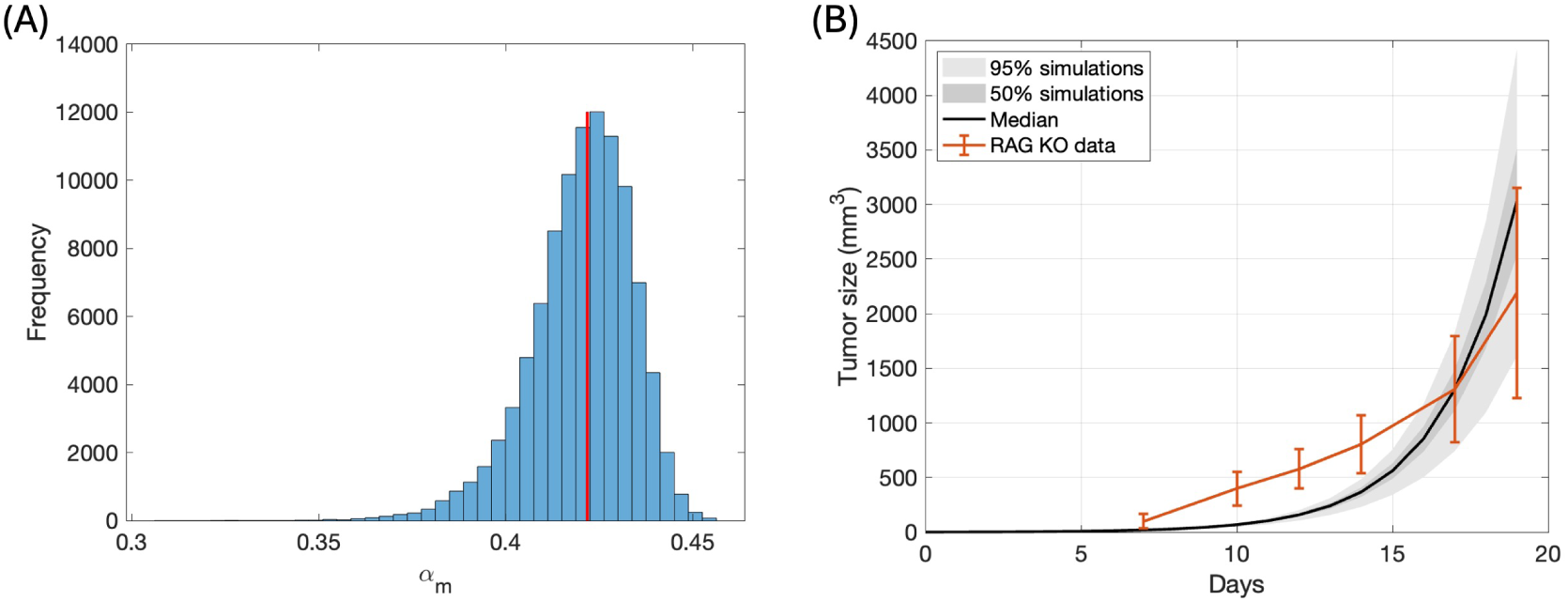
Data integration and practical identifiability analysis: distribution of tumor proliferation rate (*α_m_*) and simulated tumor growths versus experimental data in the RAG1 KO cohort.

**Figure S3:**
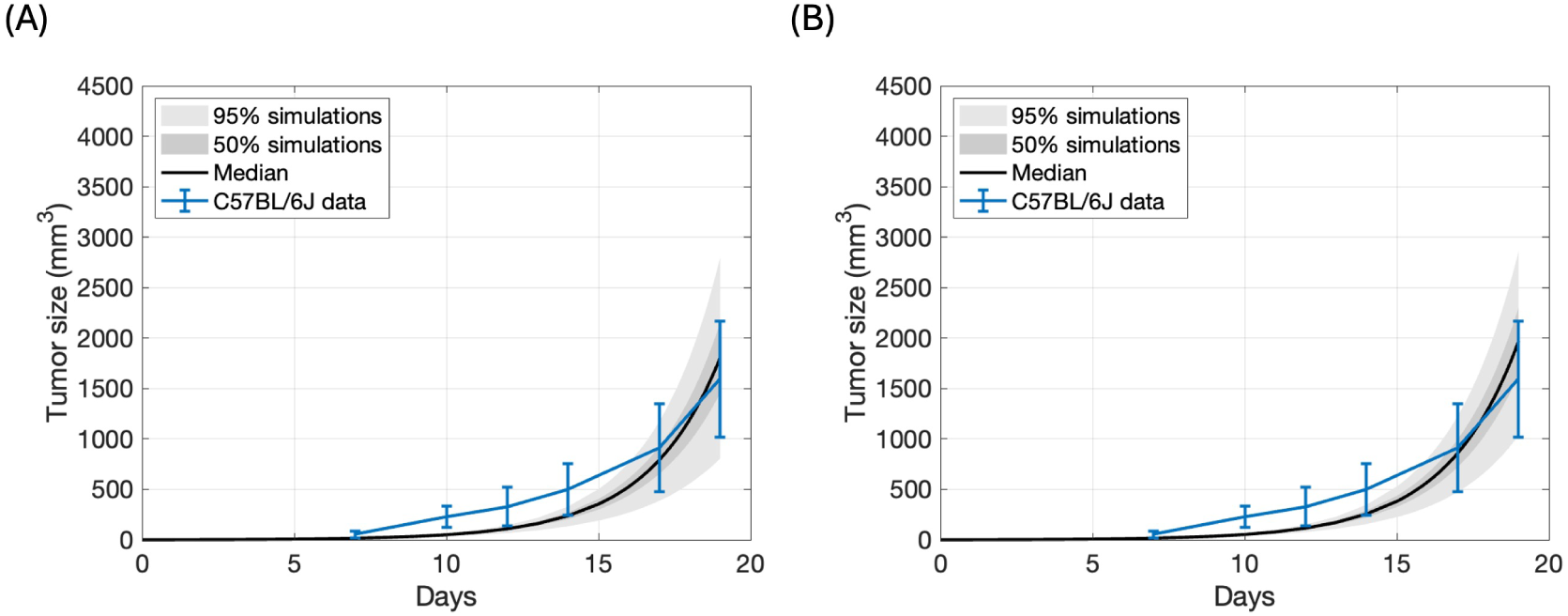
Data integration and practical identifiability analysis: indistinguishable simulated tumor growths in the virtual cohorts constructed with unidentifiable immune parameters and identifiable immune parameters, respectively. (A) Virtual cohort constructed using tumor measurements in C57BL/6J data. The posterior distributions which the cohort sampled from correspond to Figure 4A. Immune parameters *µ* and *p*_fast_ are unidentifiable. (B) Virtual cohort constructed using both RAG1 KO and C57BL/6J data. The posterior distributions correspond to Figure 4B. Immune parameters *µ* and *p*_fast_ are identifiable as a pair.

**Figure S4:**
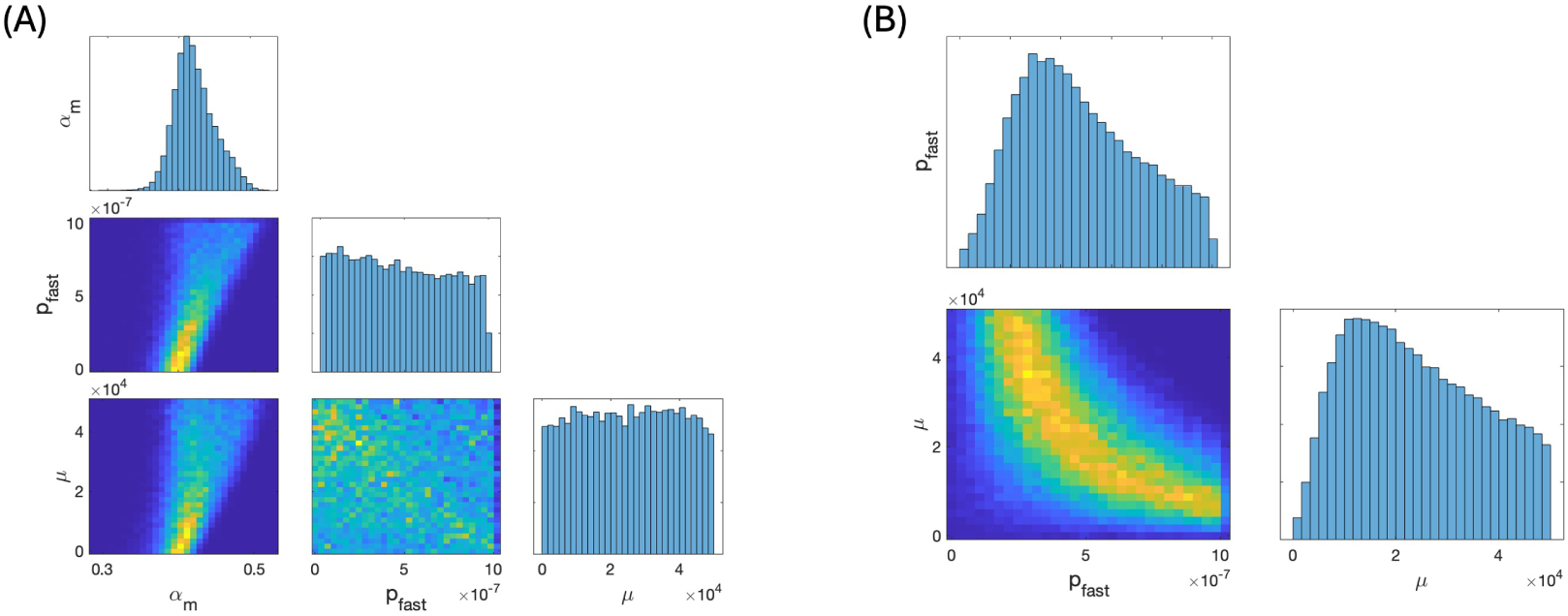
Data integration and practical identifiability analysis: posterior distributions of model parameters following practical identifiability analysis using RAG1 KO data and C57BL/6J data with endpoint CTL density measurements. The results are very similar to Figure 4.

## Notes

### Competing Interest Statement

The authors have declared no competing interest.

### Funding

- NIH/NCI: U01CA243075, K08, CA234392

